# Pairwise correlations of spiking activity changes along the ventral visual cortical pathway of macaque monkeys

**DOI:** 10.1101/220301

**Authors:** Hiroshi Tamura

**Affiliations:** Graduate School of Frontier Biosciences, Osaka University, Suita, Osaka 565-0871, Japan; Center for Information and Neural Networks, Suita, Osaka 565-0871, Japan

**Author notes:** Corresponding author: Dr. Hiroshi Tamura, Laboratory for Cognitive Neuroscience, Graduate School of Frontier Biosciences, Osaka University 2A1, Center for Information and Neural Networks, 1-4 Yamadaoka, Suita, Osaka 565-0871, Japan.

## Abstract

Spiking activity often correlates across neurons in the cerebral cortex. Precise spike-timing correlation at millisecond scale reflects underlying neural circuitries. Correlations of stimulus evoked spiking activities affects the distributions of spike counts in a multi-neuron space and reflects stimulus encoding ability of neuron populations. The present study compared the degree of spiking-activity correlation among three areas of the macaque visual system to determine whether neural circuitry and the way to encode stimulus-related information by neuron populations are unique to each area. Spiking activity of multiple single neurons was recorded in macaque primary visual cortex (V1), visual association cortex (V4), and inferior temporal cortex (IT). Cross-correlation of spike trains revealed that single-neuron pairs in IT exhibited the highest incidence of precisely correlated firing, whereas V1 pairs showed the lowest incidence. Although noise correlation, which quantifies similarity in trial-to-trial response fluctuations, differed among cortical areas, signal correlation, which quantifies similarity in stimulus preferences, did not. The degree of temporally precise correlation was positively related to the signal-correlation strength in all three cortical areas and the relation of IT pairs was stronger than that of V1 pairs. Temporally precise correlations between single-neuron and population-level activity also differed among cortical areas and V1 neurons exhibited the lowest incidence of correlated firing with population activities. The differences in spiking-activity correlations among visual cortical areas suggest that each cortical area has unique neuronal circuitry for representing information, and which performs unique neuronal computations.

**Significance Statement:** Although hierarchical information processing plays a crucial role in the brain and in artificial neural networks, whether hierarchically connected brain areas share neural circuitry and information representation remains unclear. Here, we examined spiking activity across the ventral visual system in macaque monkeys and found that cortical areas differed in how correlated their spiking activity was, suggesting that neural circuitry and the way to encode stimulus-related information by neuron populations are unique to each cortical area.

## Introduction

Spiking activity in the cerebral cortex correlates across individual neurons. Precise spiking-activity correlation at the millisecond scale reflects the organization of underlying neural circuitry (Perkel et al., 1967; Moore et al., 1970; Bryant et al., 1973). For example, correlated activity without time delay reflects common inputs to paired neurons. Precise spiking-activity correlation have been proposed to encode stimulus related information (Singer and Gray, 1995; Diesmann et al., 1999; Hirabayashi and Miyashita, 2005; Panzeri et al., 2015; Zandvakili and Kohn, 2015). Thus, studying correlations at the millisecond scale allows us to infer neural circuitry between paired neurons and information encoding by synchrony. Two other types of spike correlation related to amount of information encoded by neuron populations (Averbeck et al. 2006; Cohen and Kohn, 2011; Panzeri et al., 2015; Franke et al., 2016). Signal correlation refers to the correlation between trial-averaged spike counts of paired neurons, which reflects the similarity in stimulus-evoked responses (Gawne and Richmond, 1993; Panzeri et al., 2015). If neurons in a population encode independent information, signal correlation is low, whereas if they encode redundant information, signal correlation is high. Responses of cortical neurons varies from one trial to another. Noise correlation—or spike count correlation—refers to the correlation between the deviations of single-trial spike counts from the trial-averaged response of paired neurons, and reflects the similarity of trial-to-trial response fluctuations (Zohary et al., 1994). Relationship between degrees of noise correlation and information encoding ability of a neuron population may not be straightforward. Correlated noise may limit or enhance the ability of a neuron population to encode information depending on the sign of signal correlation (Zohary et al., 1994; Cohen and Kohn, 2011; Panzeri et al., 2015).

Such correlations in spiking activity have been examined extensively, especially in the visual cortical areas of primates (Krüger and Aiple, 1988; Ts’o and Gilbert, 1988; Gochin et al., 1991; Kreiter and Singer, 1992; Gawne and Richmond, 1993; Zohary et al., 1994; Gawne et al., 1996; Tamura et al., 1996; Roe and Ts’o, 1999; Maldonado et al., 2000; Bair et al., 2001; Tamura et al., 2004; Aggelopoulos et al., 2005; Hirabayashi and Miyashita, 2005; Kohn and Smith, 2005; Tamura et al., 2005; Kreiman et al., 2006; Smith and Kohn, 2008; Cohen and Maunsell, 2009; Kotake et al., 2009; Mitchell et al., 2009; Sato et al., 2009; Ecker et al., 2010; Huang and Lisberger, 2013; Smith and Sommer, 2013; Ecker et al., 2014; Goris et al., 2014; Lin et al., 2014; Tamura et al., 2014; Hung et al., 2015; Okun et al., 2015; see Cohen and Kohn, 2011, for a review of noise correlation). If neural circuitry and information encoding are shared among cortical areas, the degree to which spiking activity is correlated might be similar across cortical areas. Conversely, if the circuitry and information encoding are unique to each cortical area, spiking-activity correlation might differ among them. Although a few studies have directly compared correlations in spiking activity between two visual cortical areas (Gawne et al., 1996; Tamura et al., 1996; Smith and Sommer, 2013), whether spike-correlation systematically differs along hierarchically connected cortical regions remains unclear.

The present study aimed to clarify whether a variety of correlations between the spiking activity of paired neurons differ among the primary‐ (V1), middle‐ (V4), and late-stage (inferior temporal cortex, IT) visual cortical areas of macaque monkeys. Because these areas comprise the hierarchically organized ventral visual cortical pathway, they are ideal for assessing whether correlations change systematically along the cortical hierarchy. Correlations between the spiking activities of paired neurons in each cortical area revealed that some types of correlations differed among the three regions and changed systematically along the visual cortical hierarchy. Relationships between the spiking activity of single neurons and that of adjacent neurons as a population also differed among the three cortical areas.

## Materials and Methods

The general experimental procedures have been described previously (Tamura et al., 2016), and the present study analyzes the data obtained in the previous study. Briefly, neuronal activity was recorded from three monkeys *(Macaca fuscata;* body weight, 5.9-8.6 kg; Monkeys A, B, C). Recordings from IT cortex were obtained from all three monkeys and those from V4 and V1 were obtained from two monkeys (Monkeys A and C). The monkeys were anesthetized and paralyzed because body movements affect recording stability and eye movements affect the response variance and correlation of neuronal spiking activity (Gur et al., 1997; Maldonado et al., 2008; McFarland et al., 2016). Because anesthesia has been shown to increase noise correlation (Ecker et al., 2014), the correlation measures should be interpreted with caution (but see Discussion). All experiments were performed in accordance with the guidelines from the National Institute of Health (1996) and the Japan Neuroscience Society, and were approved by the Osaka University Animal Experiment Committee.

### Initial preparatory surgery

To prepare the monkeys for the recordings, a head restraint was implanted and the lateral or occipital part of the skull (over the recording region) was covered with acrylic resin. Monkeys were fully anesthetized during surgery by inhalation of 1%-3% isoflurane (Forane, Abbott Japan, Tokyo, Japan) in nitrous oxide (70% N_2_O, 30% O_2_) through an intratracheal cannula. Antibiotics (Pentcilin, Toyama Chemical, Tokyo, Japan; 40 mg/kg, i.m.) and an anti-inflammatory and analgesic agent (Voltaren, Novartis, Tokyo, Japan; or Ketoprofen, Nissin Pharmaceutical, Yamagata, Japan) were administered immediately after the surgery and during the first postoperative week. After 1-2 weeks of recovery, the monkeys’ eyes were examined to select appropriate contact lenses that would focus images placed 57 cm from the cornea on the retina. The retinal fundus was photographed to determine the position of the fovea.

### Preparation of animals for neural recording

On the day of neural recording, monkeys were sedated via intramuscular injection of atropine sulfate (0.1 mg/kg) and ketamine hydrochloride (12 mg/kg). The monkeys were then anesthetized by inhalation of 1%−3% isoflurane in nitrous oxide (70% N_2_O, 30% O_2_) through an intratracheal cannula and by an infusion of the opioid fentanyl citrate (Fentanest, Daiichi Sankyo; 0.035 mg/kg/h) in lactated Ringer’s solution. A small hole (~7 mm) was drilled in the resin-covered skull and a small slit (2 mm) was made in the dura to insert electrodes. The pupil was dilated and the lens was relaxed using 0.5% tropicamide/0.5% phenylephrine hydrochloride (Mydrin-P, Santen, Osaka, Japan). The cornea was then covered with the contact lens that included an artificial pupil (3 mm diameter). After inserting the recording electrode, vecuronium bromide (Masculax, MSD, Tokyo, Japan; 0.06 mg/kg/h) was added to the infusion solution to prevent eye movements while recording neuronal activity. After each recording session, monkeys were given analgesics and antibiotics and returned to their home cages. Each recording session lasted up to 7 h, and consecutive recording sessions in the same monkey were separated by at least one week.

### Neuronal-activity recordings

Spiking activity from multiple-single neurons was recorded in IT, V4, and V1 using a single-shaft electrode with 32 recording probes arranged linearly (A1X32-10 mm 50-413; NeuroNexus, Ann Arbor, MI, USA). The distance between the centers of adjacent recording probes was 50 µm. Because spiking activity from the same neuron could be picked up by two or more adjacent probes, single neuron activity was isolated offline using custom-made software (see Kaneko et al. 1999; Kaneko et al. 2007; Tamura et al. 2014, for details). Spikes were classified with reference to the signals recorded from five consecutive probes *(i.e.,* within 0.2 mm). Care was taken not to assign single spikes to two or more neurons. A neuron’s position along the electrode was defined as the position of the probe that recorded its spikes at the largest amplitude.

Recordings were made in 10 recording sessions for the IT, and 8 sessions each for V4 and V1. The recording sites in IT were located between the superior temporal sulcus and the anterior middle temporal sulcus, and anterior to the posterior middle temporal sulcus. Those in V4 were located between the superior temporal sulcus and the lunate sulcus, and those in V1 were located on the surface of the occipital cortex away from the lunate sulcus.

### Visual stimuli

The stimulus set comprised 64 color images (8 each from 8 categories of natural objects: stones, trees bark, leaves, flowers, fruits and vegetables, butterfly wings, feathers, and skin and fur) and a blank image with the same pixel values as the background (Tamura et al., 2016). All images were 6° in visual angle. Images were presented on a liquid crystal display (CG275W, Eizo, Ishikawa, Japan), which was regularly calibrated with the internal calibrator and checked with a spectrometer (Minolta CS-1000, Tokyo, Japan). The luminance values of the white and black areas of the display were 125 cd/m^2^ and 1.3 cd/m^2^, respectively. Each image was presented for 200 ms with the inter-stimulus interval of 200 ms. Stimulus onsets and offsets were recorded using a photodiode attached to the monitor. Each stimulus image was presented 25 times during recording. Images were presented to the eye contralateral to the recording hemisphere, while the other eye remained closed. For IT neurons, images were presented to the fovea because the receptive fields (RFs) of IT neurons include the fovea and respond well to stimuli located there (Gross et al., 1972). For V4 and V1 neurons, images were presented to the center of the RF, as determined by listening to the amplified multiunit activity while visually stimulating the neurons with hand-held circular disks, bars, or grating patterns.

### Data analysis

The magnitude of visually evoked responses were calculated for each stimulus by subtracting the spontaneous firing rate during the 100-ms period immediately preceding stimulus onset from the raw firing rate during the 200-ms stimulus presentation. The window for the 200-ms stimulus-presentation period was shifted to 80 ms after stimulus onset for IT neurons to compensate for response latency. The responsiveness of each neuron was evaluated by comparing the firing rates during the stimulus presentation period across stimuli (*P* < 0.01, Kruskal-Wallis test).

Correlations between paired neurons were measured in three ways. Precise spiking-activity correlations at millisecond-scale were calculated by cross-correlating paired spike trains at a temporal resolution of 1 ms and constructing cross-correlograms (CCGs). Raw CCGs were constructed using all spikes collected from the paired neurons during the entire recording, including both visual-stimulation and inter-stimulus periods. The spike count in each CCG bin was normalized with respect to the geometric mean of the spike counts of paired neurons. Because visual stimulation activated the two neurons of a pair almost simultaneously, a raw CCG reflects both the correlation due to the stimulus-locked activation (stimulus coordination) and that due to functional connections between the two neurons (neuronal correlation). To separate these contributions, a shift-predicted CCG was calculated by shifting the timing of the spike trains from one of the two neurons by one trial (Perkel et al. 1967; Toyama et al. 1981; see Tamura et al., 2014 for details). The center peaks of the CCGs were detected by comparing the spike counts in each 1 −ms bin of a raw CCG with those from the corresponding shift-predicted CCG bin (*P* < 0.0001, binomial test). The comparison was performed within a time window of ± 10.5 ms to detect significant peaks at or around the 0-ms bin (center peaks). The shift-predicted CCG was subtracted from the raw CCG to obtain the shift-corrected CCG. The height and width of the center peaks of shift-corrected CCGs were then quantified. The heights were log transformed when performing statistical analyses. Because spikes were sorted using the signals recorded from five consecutive probes and because the method cannot separate spikes that occur within 0.2 ms, the spike counts in the 0-ms bin (± 0.5 ms) of the raw CCG for paired neurons recorded from the same or nearby probes (< five probes, 0.2 mm) might be underestimated. In other words, those separated by more than five consecutive probes were not affected by this problem.

Signal correlation quantifies the similarity between the trial-averaged responses (signal) of a pair of neurons to the 64 stimuli. Signal correlation was assessed using the Pearson correlation coefficient between the two sets of trial-averaged responses. Noise correlation quantifies the similarities in single-trial response deviations (noise) from the signal between paired neurons. The noise for each stimulus was calculated by subtracting the signal from single-trial responses, then dividing the result by the standard deviation across trials (Zohary et al., 1994). Noise correlation was assessed using the Pearson correlation coefficient between the two sets of single-trial response deviations. Signal correlations and noise correlations were Fisher transformed when performing statistical tests. Visually responsive neurons (*P* < 0.01, Kruskal-Wallis test) were selected for the signal correlation and noise correlation analyses.

To analyze the activity of neuronal populations, population spike trains were constructed by summing the spike trains from all other simultaneously recorded neurons with 1-ms bins (see Fig. 4A). If two or more neurons emit spikes in a 1-ms bin, the bin takes a value larger than one; for example, if three neurons fire in a bin, the value of the bin is three. To quantify correlation between the activity of single neurons and that of neuronal populations at the millisecond scale, spike trains of single neurons were cross-correlated with population spike trains at a temporal resolution of 1 ms. The resultant CCGs are termed population-neuron CCGs (Okun et al., 2015, refer to them as spike-triggered population rates). Population-neuron signal correlation was calculated using the Pearson correlation coefficient between the trial-averaged responses of a single neuron and those of the summed responses of all other simultaneously recorded neurons. Population-neuron noise correlation was assessed using the Pearson correlation coefficient between a single-trial response deviation of a single neuron and that of the summed responses of all other simultaneously recorded neurons.

### Experimental design and statistical analysis

For statistical analyses, all the data were pooled. The details of number of neurons and neuron pairs were listed in the Results section. Statistical threshold (*P* value) was set at 0.01, except for CCG-peak detection, where it was set at 0.0001. Effect sizes for non-parametric data (r) were calculated with the formula

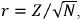

where *Z* and *N* represent the z-transformed Mann-Whitney U-test statistic and the number of samples, respectively. Effect sizes for chi-square tests (Cramer’s V) were calculated with the formula

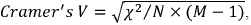

where *χ*^2^ is the chi-square statistic, *N* is the number of samples, and *M* is the smaller of the two values among the number of the rows and columns of the contingency table. The threshold for the effect size was set at 0.1. Thus, differences among populations were regarded as significant if the *P* value was less than 0.01 and the effect size was larger than 0.1 (Cohen, 1988). Statistical testing was performed using MATLAB (The MathWorks, MA) and provided exact p-values except for the test for the similarity of two independent correlation coefficients that was performed using a statistical software R (https://www.r-project.org) with the ‘psych’ package, which does not return the exact p-values if p-values are smaller than 0.01.

## Results

Neuronal activity was recorded from 189, 149, and 171 neurons from IT, V4, and V1, respectively. From these neurons, 2,082 IT neuron pairs, 1,797 V4 pairs and 2,261 V1 pairs were constructed. Note that single neurons contributed to multiple pairs if they were recorded simultaneously.

### Comparisons of cross-correlated spiking activity among IT, V4, and V1 neuron pairs

Correlations between the spiking activities of single-neuron pairs at the millisecond scale were compared among IT, V4, and V1. The cross-correlation analyses of the spike trains from paired neurons provided CCGs at a temporal resolution of 1 ms. The presence/absence of CCG center peaks was judged statistically by comparing the spike counts in each 1-ms bin of raw CCGs with those of shift-predicted CCGs within ± 10.5 ms (*P* < 0.0001, binomial test). The center peak incidence for the CCGs was 14% for IT pairs (298/2,082), 10% for V4 pairs (186/1,797), and 5% for V1 pairs (103/2,261; Fig. 1A). These incidences differed across the three regions (*P* = 2.2×10^-1^, effect size = 0.14, chi-square test). Thus, each cortical area had a unique degree of spiking-activity correlation at the millisecond scale, and the incidence of correlated neuron pairs changed systematically along the cortical hierarchy.

Width of CCG peaks also differed systematically along the cortical hierarchy. Visual inspection showed broad peaks in the CCGs of IT pairs and sharp peaks in those of V1 pairs (Fig. 1B-D). The peak width of the average shift-corrected CCGs across neuron pairs with significant center peaks (298 IT pairs, 186 V4 pairs, and 103 V1 pairs) can quantify this impression by fitting the exponential function,

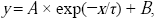

where *y* and *x* are the normalized spike count and CCG time delay, respectively; *A* is amplitude and *B* is offset; and **τ** is a time constant that corresponds to the time delay until the amplitude decreases to 37% of the peak. The time constant (*τ*) was used as a measure of the peak width, and was 40.1 ms for IT pairs (*R* ^2^ = 0.95), 21.6 ms for V4 pairs (*R* ^2^ = 0.85), and 13.4 ms for V1 pairs (*R* ^2^ = 0.60). Thus, as the initial impression indicated, peak width was relatively broad for IT pairs and sharp for V1 pairs. Although fitting an exponential function to the average CCGs was successful, doing so for individual pairs was only successful for a small number of pairs. Therefore, a *width index* (which Tamura et al., 2014, refers to as the peak-to-area ratio) was used as another measure of CCG width. *Width index = area under the curve/peak height,* where the *area under the curve* is the sum of the spike counts under the shift-corrected CCGs, and *peak height* is the spike count at the CCG center peaks. Pairs with a negative area under the curve (18 IT pairs, 20 V4 pairs, 17 V1 pairs) were excluded from this analysis. The value of the index ranged from 1 for neuron pairs that showed the sharpest peak (one bin width) up to 500 (the number of bins in CCGs) for neuron pairs that showed much broader peaks. IT pairs had the largest width index with a median of 30.9 (280 pairs), followed by V4 pairs with a median of 18.7 (166 pairs), and V1 pairs with the smallest median of 7.9 (86 pairs; Fig. 1E). The width index differed among the three cortical areas (*P* = 3.0×10^-21^, Kruskal-Wallis test; effect size, 0.24 for IT-V4, 0.48 for IT-V1, 0.35 for V4-V1). These results suggest that the hierarchical order in the ventral visual pathway is reflected in the temporal properties of spike interactions.

IT pairs exhibited a higher incidence of CCG peaks and broader peak width than V1 pairs. The higher incidence of CCG peaks among IT pairs was likely due to the higher incidence of CCGs with broad peaks. To examine this possibility, the incidences of CCGs with sharp peaks and broad peaks were examined separately among cortical areas. Sharp/broad peaks were arbitrarily defined as peaks with a width index smaller/larger than the median of the V1 pairs (width index, 7.9). The incidence of CCGs with sharp peaks was low and did not differ among the IT (1.6%, 34/2,082), V4 (2.1%, 38/1,797), and V1 (1.9%, 43/2,261; *P* = 0.540, effect size = 0.01, chi-square test), whereas that with broad-peaks did (IT: 11.8%, 246/2,082; V4: 7.1%, 128/1,797; V1: 1.9%, 43/2,261; *P* = 2.3×10^-37^, effect size = 0.17, chi-square test). These results suggest that the higher incidence of CCG peaks in IT pairs is likely to derive from higher incidences of CCGs with broader peaks.

**Figure 1.**
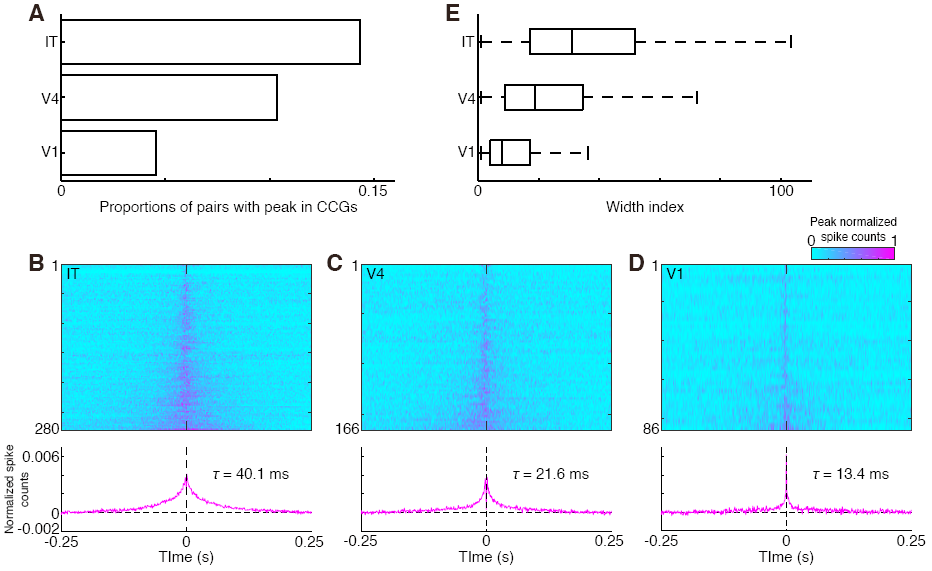
Cross-correlation analysis of spiking activity for paired neurons recorded from IT, V4, and V1. ***A**,* Comparison of the proportions of pairs with significant CCGs center-peaks among IT (n = 2,082), V4 (n = 1,797), and V1 (n = 2,261). ***B-D**,* Shift-corrected CCGs with significant center peaks for IT (B, top, n = 280), V4 (C, top, n = 166), and V1 (D, top, n = 86). Each line in a color plot corresponds to individual CCG, which was normalized with its CCG-peak amplitude and color coded. Color coded CCGs were sorted according to width index. Average shift-corrected CCGs across the CCGs of pairs with significant center-peaks are shown for IT (B, bottom), V4 (C, bottom), and V1(D, bottom). The horizontal dashed line is the baseline and the vertical dashed line is the zero-time delay. The time constant (r) that corresponds to the time delay until the amplitude decreases to 37% of the peak is provided in the plots. ***E**,* Comparison of the width index of CCG-center peak among IT, V4, and V1. The index takes a value of one for neuron pairs with sharp peaks one-bin in width and takes larger values for neuron pairs with much broader peaks. The center of each box plot is the median, and the right and left of the box are the upper and lower quartiles, respectively. Attached whiskers connect the most extreme values within 150% of the interquartile range from the end of each box.

### Comparisons of signal and noise correlations among IT, V4, and V1 neuron pairs

Signal and noise correlations across the three brain regions were used to examine the correlations between spiking activities over a longer timescale. Signal correlation is the correlation of trial-averaged responses (signal) during the stimulus presentation period (200 ms), and noise correlation is the correlation of the single-trial response deviation (noise) from the signal. Visually responsive neurons (*P* < 0.01, Kruskal–Wallis test; 141 IT neurons, 86 V4 neurons, and 107 V1 neurons) were used for the analysis, yielding 1,291 IT pairs, 759 V4 pairs, and 1,014 V1 pairs.

The median signal correlation was positive in all the three cortical areas (IT, 0.070; V4, 0.039; V1, 0.083; Fig. 2A, B). The median signal correlation across IT pairs was different from zero (*P* = 1.1×10^−39^, effect size = 0.37, Wilcoxon signed-rank test; Fig. 2A) and the signal correlation of IT pairs was different from that between IT neurons recorded from different sites (median = 0.018, *P* = 4.1×10^−28^, effect size = 0.11, Mann-Whitney U-test). In both V4 and VI, the median signal correlation was different from zero (V4, *P* = 2.2×l0^−12^, effect size = 0.25; VI, *P* = 6.7×l0^−32^, effect size = 0.37; Fig. 2B), but their signal correlation was not different from the signal correlation between neurons recorded from different sites (V4, median = 0.026, *P* = 2.6×l0^−4^, effect size = 0.06; V1, median = 0.065, *P* = 6.5×l0^−4^, effect size = 0.05). The signal correlations did not differ among IT, V4, and V1 (*P* = 0.0014, Kruskal–Wallis test; effect size, 0.06 for IT-V4, 0.02 for IT-V1, 0.08 for V4-V1).

The median noise correlations were also positive (IT, 0.020; V4, 0.015; VI, 0.007; Fig. 2C, D), and was different from zero (IT, *P* = 1.4×10^−62^, effect size = 0.46; V4, *P* = 3.7×l0^−23^, effect size = 0.36; V1, *P* = 1.1×10^−11^, effect size = 0.21, Wilcoxon signed-rank test) in all three cortical areas. Consistent with the differences in the incidences of CCG peaks among cortical areas, the noise correlations differed between IT and V1 (*P* = 2.1×10^−16^, Kruskal–Wallis test; effect size, 0.09 for IT-V4, 0.17 for IT-V1, 0.09 for V4-V1), and IT pairs had the highest noise correlation and VI pairs had the lowest noise correlation.

**Figure 2.**
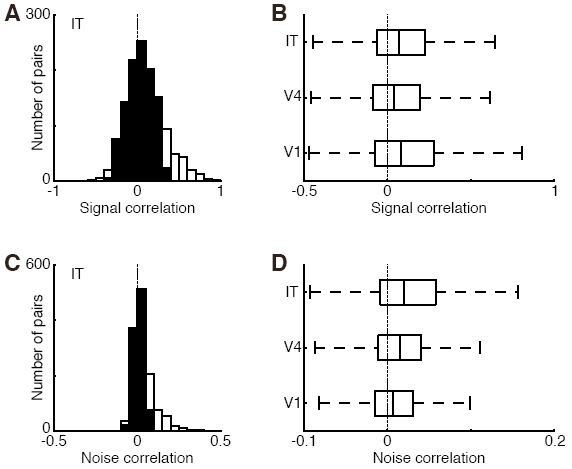
Signal correlation and noise correlation of paired neurons recorded from IT, V4, and V1. ***A**,* Frequency distribution of signal correlation for IT pairs. Open column, significant correlation (*P* < 0.01, test of independence). Filled column, non-significant correlation. Vertical dashed line, zero signal correlation. ***B**,* Comparison of signal correlation across IT, V4, and V1. Vertical dashed line, zero signal correlation. ***C**,* Frequency distribution of noise correlation for IT pairs. Other conventions are as in A. ***D**,* Comparison of noise correlation across IT, V4, and V1. Other conventions are as in B.

### Relationship between CCG-peak height and signal correlation or noise correlation

Analysis revealed that CCG-peak height correlated positively with signal correlation in all three cortical areas, (IT, *r* = 0.48, *P* = 8.7×10^-74^; V4, *r* = 0.42, *P* = 1.8×10^-33^; V1 *r* = 0.28, *P* = 3.6×10^-19^; test for independence; Fig. 3A-C), meaning that if two neurons received common inputs, they tended to share stimulus preferences. However, the degree of correlation differed depending on the region. V1 pairs exhibited lower correlation than pairs in other areas (V1-IT, *P* < 0.01; V1-V4, *P* < 0.01; V4-IT, *P* = 0.12, test for the similarity of two independent correlation coefficients; Fig. 3D), meaning that the relationship between the organization of common inputs and the similarity in stimulus preferences are less clear in V1. These results can explain the finding that signal correlation did not differ among cortical areas, despite the differences in the incidences of CCG peaks among cortical areas.

Similar to signal correlation, CCG-peak height correlated positively with noise correlation in all three cortical areas (IT, *r* = 0.69, *P* = 1.4×10^-183^; V4, *r* = 0.51, *P* = 3.4×10^-51^; V1 *r* = 0.40, *P* = 7.5×10^-41^; test for independence; Fig. 3E-G). Again, the degree of correlation depended on the cortical areas. IT neurons exhibited higher positive correlation than pairs in other areas, and like with signal correlation, V1 neurons displayed lower positive correlation than pairs in other areas (IT-V4, *P* < 0.01; IT-V1, *P* < 0.01; V4-V1, *P* = 0.01; Fig. 3H). These results show that the relationship between the organization of common inputs and the similarity in trial-to-trial response fluctuations differed among cortical areas.

**Figure 3.**
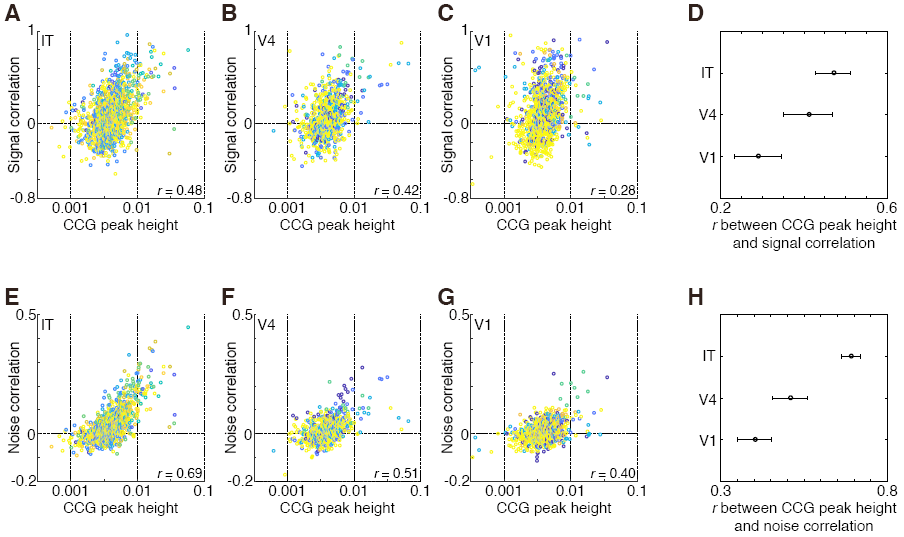
Relationships between CCG-peak height and signal correlation or noise correlation. ***A-C**,* Relationship between CCG-peak height and signal correlation in IT (A), V4 (B), and V1 (C) pairs. CCG-peak heights are plotted on a logarithmic scale. Each point represents one neuron pair. Each color represents pairs obtained from a single recording session. The correlation coefficient (*r*) between CCG-peak height and signal correlation is provided in the plots. ***D**,* Comparison of the correlation coefficient (*r*) between CCG-peak height and signal correlation across IT, V4 and V1. Attached bars are 95% confidence intervals. ***E-G**,* Relationship between CCG-peak height and noise correlation in IT (E), V4 (F), and V1 (G) pairs. Other conventions are as in A-C. ***H**,* Comparison of correlation coefficient (*r*) between CCG-peak height and noise correlation across IT, V4, and V1. Other conventions are as in D.

### Relationships between single-neuron and population-neuron activities

Although significant correlations between the spiking-activities of paired single neurons were observed in all three cortical areas, they were generally weak; *i.e.,* low incidences of CCG center peaks and positive but near-zero signal and noise correlations. Such weak single-neuron correlations suggest that most neurons emit spikes independently from adjacent neurons. However, weak single-neuron correlations may lead to much stronger correlations between single-neuron activity and that of adjacent neurons as a population. The correlations between single-neuron and population-neuron activities were calculated to examine this issue. For each single-population correlation, the population-neuron activity was constructed from the activity of all the other simultaneously recorded neurons (see Methods for details; Fig. 4A). Population-neuron CCGs were generated by cross-correlating single-neuron spike trains with population-neuron spike trains that were obtained by simply summing the spike trains of all the other simultaneously recorded neurons. The center-peak incidences in the population-neuron CCGs were 79% for IT neurons (149/189), 80% for V4 neurons (119/149), and 58% for V1 neurons (99/171; Fig. 4B). These incidences were significantly higher than those observed in single-neuron CCGs (IT, *P* = 3.1×10^-15^, effect size = 0.45; V4, *P* = 1.6×10^-76^, effect size = 0.51; V1, *P* = 1.2×10^-71^, effect size = 0.49, chi-square test; see Fig. 1A). These results suggest that the relationship between single-neuron spiking activity and the activity of adjacent neurons as a population is stronger than that between pairs of single neurons. The center-peak incidence in population-neuron CCGs differed across the three cortical areas (*P* = 2.4×10^-6^, effect size = 0.43, chi-square test), indicating that the relationship between single-neuron and population-neuron activities differed among cortical areas; *i.e.,* the correlations between single-neuron and population-neuron spiking activities were weaker in V1 than in V4 or IT.

**Figure 4.**
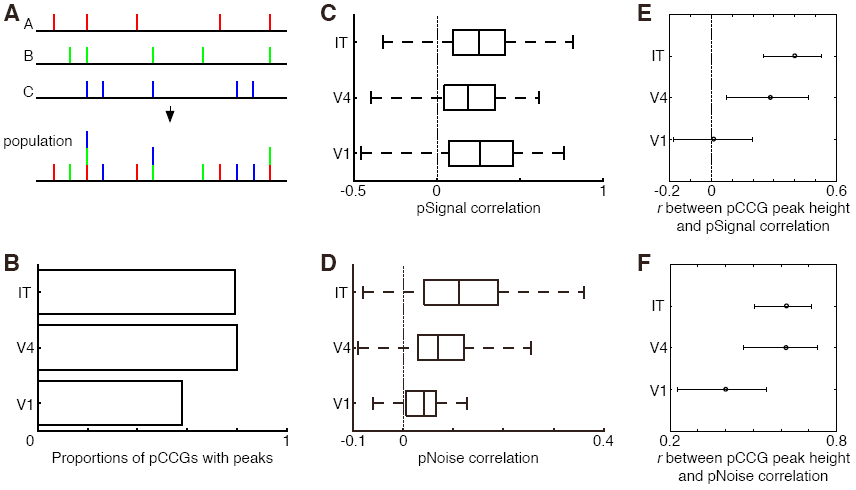
Relationships between single-neuron and population-neuron activities. ***A**,* Construction of population spike trains (population) from spike trains of three neurons (A, B and C). ***B**,* Comparison of proportions of neurons with significant (*P* < 0.0001, binomial test) center peaks in population CCGs (pCCGs) for IT (n = 189), V4 (n = 149), and V1 (n = 171). ***Q*** Comparison of population signal correlation (pSignal correlation) for IT, V4, and V1. Vertical dashed line, zero correlation. ***D**,* Comparison of population noise correlation (pNoise correlation) across IT, V4, and V1. Other conventions are as in C. ***E**,* Comparison of correlation coefficient (*r*) between population CCG-peak height and population signal correlation across IT, V4, and V1. Attached bars are 95% confidence interval. Vertical dashed line, zero correlation. ***F**,* Comparison of correlation coefficient (*r*) between population CCG-peak height and population noise correlation across IT, V4, and V1. Other conventions are as in E.

**Figure 5.**
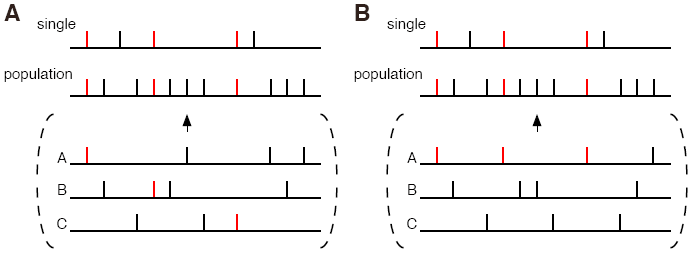
Two possible scenarios that explain the higher incidences of peaks in CCGs for single/population pairs than for single/single pairs. Here, the spike train of a single neuron (single) was correlated with that of the neuron population (population) derived from neuron A-C. ***A**,* One possible scenario (democratic) is that all neurons contribute to the correlated firings, but their contributions are small. Vertical bars represent timings of spikes. Red and black bars represent correlated and independent spikes, respectively. Correlated spikes (red vertical bars) are observed in all the neurons. ***B**,* The other scenario (despotic) is that only one or a few neurons (here, neuron A) contribute to the correlated activity. Correlated spikes (red vertical bars) are observed only in neuron A. Other conventions are as in A.

Consistent with the higher incidences of peaks in population-neuron CCGs than in single-neuron CCGs, population-neuron signal and noise correlations were stronger than their single-neuron counterparts. Population-neuron signal correlation was 0.25 (median) for IT (n = 141), 0.19 for V4 (n = 86), and 0.26 for V1 (n = 107; Fig. 4C). Population-neuron signal correlation was stronger than single-neuron signal correlations in all three cortical areas (IT, *P* = 3.3×10^-14^, effect size = 0.20; V4, *P* = 3.6×10^-7^, effect size = 0.18; V1, *P* = 1.4×10^-8^, effect size = 0.17, Mann-Whitney U-test; see Fig. 2B). Population-neuron noise correlation was 0.11 (median) for IT (n = 141), 0.07 for V4 (n = 86), and 0.04 for V1 (n = 107; Fig. 4D), and was higher than single-neuron noise correlation in all three cortical areas (IT, *P* = 2.4×10^-25^, effect size = 0.27; V4, *P* = 1.5×10^-15^, effect size = 0.27; V1, *P* = 7.0×10^-12^, effect size = 0.20; Mann-Whitney U-test; see Fig. 2D). These results suggest that single-neuron responses are much more similar to the averaged responses of adjacent neurons than to the responses of adjacent single neurons. Population-neuron signal correlation did not differ among the three regions (*P* = 0.051, Kruskal-Wallis test; effect size, 0.14 for IT-V4, 0.02 for IT-V1, 0.16 for V4-V1), whereas population-neuron noise correlation did (*P* = 8.9×10^-10^, Kruskal-Wallis test; effect size, 0.18 for IT-V4, 0.40 for IT-V1, 0.27 for V4-V1).

Next, analyses examined whether single/population pairs with higher peaks in population-neuron CCGs tended to show higher population-neuron signal and noise correlations. The peak height of population-neuron CCGs correlated positively with population-neuron signal correlation in IT and V4, but not in V1 (IT, *r* = 0.389, *P* = 1.83×10^-6^; V4, *r* = 0.288, *P* = 0.0071; V1, *r =-* 0.0041, *P* = 0.967, test for independence; Fig. 4E). V1 pairs had significantly weaker correlation than IT pairs (V1-IT, *P* < 0.01; V1-V4, *P* = 0.04; V4-IT, *P* = 0.41, test for the similarity of two independent correlation coefficients). In all three cortical areas, the peak height of population-neuron CCGs correlated positively with population-neuron noise correlation (IT, *r* = 0.617, *P* = 3.58×10^-16^; V4, *r* = 0.617, *P* = 2.57×10^-10^; V1, *r* = 0.400, *P* = 1.95×10^-5^, test for independence; Fig. 4F). The relationship between the peak height of population-neuron CCGs and population-neuron noise correlation did not differ among cortical areas (IT-V4, *P* = 0.99; IT-V1, *P* = 0.02; V4-V1, *P* = 0.04).

## Discussion

The present study aimed to clarify whether the structures of spiking-activity correlations differ among hierarchically organized visual cortical areas of macaque monkeys. Among IT, V4, and V1, IT pairs showed the highest incidences of center peaks in their CCGs, whereas V1 pairs showed the lowest. The strength of signal correlations, however, did not differ among the three cortical areas. Correlation between CCG-peak height and signal correlation was stronger in IT than in V1. Analyses of population-neuron activity revealed a similar tendency. Thus, the activity correlations between single neurons, as well as those between single neurons and the surrounding population, differed among cortical areas.

### Each cortical area has unique local neuronal circuitry

The incidence of center peaks in the CCGs differed among cortical areas, with IT pairs showing the highest incidences and V1 pairs the lowest. Three possible confounding factors that might affect the estimated incidences of center peaks were considered. One is the difference in spike counts used for cross-correlation analysis among the three cortical areas (IT, 509-19,906; V4, 521-24,332; V1, 500-11,752). Reanalysis of the incidences of neurons that emitted spikes in a limited range (500-12,000 spikes) confirmed the result, *i.e.,* center-peak incidence was highest for IT pairs (13.8%, 266/1,924), in-between for V4 pairs (10.0%, 176/1,762), and lowest for V1 pairs (4.6%, 103/2,261). A second possible confounding factor is the distance between paired neurons. However, this is a non-issue because the estimated neuron-to-neuron distances did not differ among the three cortical areas (*P* = 0.434, Kruskal-Wallis test). A third possible confounding factor is underestimation of peak incidences due to the spike-overlapping problem. To circumvent this problem, the incidences of center peaks were compared only using neurons that were separated by more than five probes, and thus did not suffer from the spike-overlapping problem. This analysis confirmed the result; *i.e.,* the center-peak incidence in CCGs were highest for IT pairs (7.5%, 105/1,399), in-between for V4 pairs (6.6%, 80/1,204), and lowest for V1 pairs (4.2%, 62/1,484). From the above analyses, the incidence of center peaks in CCGs can be concluded to differ among the three cortical areas.

Because center peaks in the CCGs can be interpreted as a sign of common input onto paired neurons (Perkel et al., 1967), the degree of input sharing can be inferred to change systematically along the ventral visual cortical pathway. The systematic changes in the incidence of CCG center peaks is consistent with the systematic changes in morphological properties along the pathway; *i.e.,* dendrites and axon terminal arbors occupy a larger area in later-stage visual cortical areas than those in the primary visual cortex (Blasdel and Lund, 1983;

Amir et al., 1993; Saleem et al., 1993; Elston et al., 1999; Tanigawa et al., 2005). These morphological properties allow IT neurons to share inputs with many other neurons. The higher degree of input sharing observed in IT cortex may contribute to generating selective responses to complex objects by integrating a variety of visual cues (Gross et al., 1972; Perrett et al., 1982; Desimone et al., 1984; Tanaka et al., 1991; Tamura and Tanaka, 2001; Tamura et al., 2016).

Despite the differences in the organization of common inputs, signal correlation did not differ among cortical areas. This result points to the possibility that the relationship between the organization of neuronal circuitry and the similarity in stimulus preferences differed among cortical areas. Indeed, correlation between CCG-peak height and signal correlation differed among cortical areas and was stronger in IT than in V1. V1 neurons share stimulus preferences without sharing inputs, whereas IT neurons share stimulus preferences precisely through their shared inputs. Broad peak widths in the CCGs of IT-neuron pairs are likely to be a basis for the stronger relationship between CCG-peak height and signal correlations.

Although incidence of CCG center peaks for paired single-neuron activity (single/single CCG peaks) in the three cortical areas was generally low (< 14%), incidence in population CCGs (single/population CCG peaks) was high (> 58%), meaning that weak pairwise single-neuron correlations led to much stronger correlations between single-neuron and population-neuron activity. The higher incidence of single/population CCG peaks than single/single CCG peaks can be explained by two alternative scenarios. One possible scenario (democratic) is that all neurons contribute to the correlated spiking activity, but each contributes only a small fraction (Fig. 5A). In this scenario, weak input sharing across neurons in a population generates peaks in population CCGs. The other scenario (despotic) is that only one or a few neurons contribute to the correlated activity (Fig. 5B). In this scenario, strong input sharing between only two or a few neurons generates peaks in population CCGs. Our preliminary analysis suggests that the first scenario (democratic) is likely because in most cases, removing spike trains one-by-one did not alter the shape of the population CCGs (IT, 149/149; V4, 115/119; V1, 98/99).

### Neuron populations in each cortical area encode information in a unique manner

Signal correlations and noise correlations obtained in the present study are similar to those obtained in previous studies from monkeys given anesthesia as well as awake monkeys (Gawne and Richmond, 1993; Gawne et al., 1996; Kohn and Smith, 2005; Tamura et al., 2005; Kreiman et al., 2006; Smith and Kohn, 2008; Cohen and Maunsell, 2009; Kotake et al., 2009; Mitchell et al., 2009; Sato et al., 2009; Ecker et al., 2010; Smith and Sommer, 2013; Ecker et al., 2014; Tamura et al., 2014; Okun et al., 2015; see Cohen and Kohn, 2011 for review). Recently, Ecker et al. (2014) reported that opioid anesthesia induced coordinated fluctuations in neuronal activity and enhanced noise correlation (see also Goris et al., 2014). However, in the present study in which animals were given fentanyl citrate, noise correlations were generally low (0.009-0.034). The different results might be related to differences in the dosage of fentanyl citrate, as pointed out by Ecker et al. (2014). Several other factors also affect the coordinated fluctuations of neural activity (Schulz et al., 2015). These confounding factors make inter-area comparisons of results from different studies difficult (Cohen and Kohn, 2011).

Because noise correlation relates to the information-encoding ability of paired neurons (Averbeck et al. 2006; Cohen and Kohn, 2011; Franke et al., 2016), differences in noise correlation across cortical regions suggest that representations of visual information by populations of local neurons changes systematically along the ventral visual pathway. Further, because synchronous activity encodes stimulus related information (Singer and Gray, 1995; Diesmann et al., 1999; Hirabayashi and Miyashita, 2005; Zandvakili and Kohn, 2015), differences in incidences in synchronous activity, *i.e.,* CCG peaks, across cortical regions suggest that representations of information via synchrony also changes systematically along the ventral visual pathway.

Differences in CCG peak-widths may relate to differences in the information integration window. Peak-widths changed systematically along the ventral visual pathway. IT pairs had the broadest CCG peaks and V1 pairs had the narrowest peaks among the three cortical areas, as expected from a theoretical study (Tanimoto et al., 2006). A longer temporal receptive window in later-stage cortical areas has been reported (Hasson et al., 2008; Honey et al., 2012). A longer integration window is beneficial for integrating a variety of visual cues, which are likely to be processed in different speeds. The differences in temporal CCG properties might derive from the differences in temporal firing properties (Gochin et al., 1991; Murray et al., 2014), number of inputs (Krüger and Aiple, 1988), types of inputs (excitatory versus inhibitory; Perkel et al., 1967), the effects of non-sensory inputs (Goris et al., 2014), or their combinations. Currently, the factors that contribute to the difference are unknown.

From the present results, each cortical area can be inferred to have unique neural circuitry and information representation. Computations or algorithms performed by neuronal circuitry have been repeatedly reported to be shared among cortical areas (Creutzfeldt, 1977; Douglas and Martin, 2004; Carandini and Heeger, 2012). However, if we assume that neural circuitry and information representation are optimized for specific computations, differences in neural circuitry and representation among cortical areas might suggest that each cortical area performs unique computations and canonical view of cortical computations should be reexamined.

## Acknowledgements

I thank H. Kaneko for technical assistance and R. Takeuchi for valuable comments on an earlier version of the manuscript. Monkeys were provided by the National BioResource Project of MEXT, Japan. This work was supported by Grant-in-Aid for Scientific Research on Innovative Areas: “Innovative SHITSUKAN Science and Technology (No. JP15H05921) and “Non-linear Neuro-oscillology” (No. JP16H01612) from MEXT, Japan. We thank Adam Phillips, PhD, from Edanz Group (www.edanzediting.com/ac) for editing a draft of this manuscript. The authors declare no competing financial interests.

